# Online Phylogenetics using Parsimony Produces Slightly Better Trees and is Dramatically More Efficient for Large SARS-CoV-2 Phylogenies than *de novo* and Maximum-Likelihood Approaches

**DOI:** 10.1101/2021.12.02.471004

**Authors:** Bryan Thornlow, Alexander Kramer, Cheng Ye, Nicola De Maio, Jakob McBroome, Angie S. Hinrichs, Robert Lanfear, Yatish Turakhia, Russell Corbett-Detig

**Affiliations:** Department of Biomolecular Engineering, University of California, Santa Cruz; Santa Cruz, CA 95064, USA; Genomics Institute, University of California, Santa Cruz; Santa Cruz, CA 95064, USA; Department of Electrical and Computer Engineering, University of California, San Diego; San Diego, CA 92093, USA; European Molecular Biology Laboratory, European Bioinformatics Institute (EMBL-EBI), Wellcome Genome Campus; Cambridge CB10 1SD, UK; Department of Ecology and Evolution, Research School of Biology, Australian National University; Canberra, ACT 2601, Australia

**Keywords:** SARS-CoV-2, phylogenetics, parsimony, maximum likelihood, optimization

## Abstract

Phylogenetics has been foundational to SARS-CoV-2 research and public health policy, assisting in genomic surveillance, contact tracing, and assessing emergence and spread of new variants. However, phylogenetic analyses of SARS-CoV-2 have often relied on tools designed for *de novo* phylogenetic inference, in which all data are collected before any analysis is performed and the phylogeny is inferred once from scratch. SARS-CoV-2 datasets do not fit this mould. There are currently over 10 million sequenced SARS-CoV-2 genomes in online databases, with tens of thousands of new genomes added every day. Continuous data collection, combined with the public health relevance of SARS-CoV-2, invites an “online” approach to phylogenetics, in which new samples are added to existing phylogenetic trees every day. The extremely dense sampling of SARS-CoV-2 genomes also invites a comparison between likelihood and parsimony approaches to phylogenetic inference. Maximum likelihood (ML) methods are more accurate when there are multiple changes at a single site on a single branch, but this accuracy comes at a large computational cost, and the dense sampling of SARS-CoV-2 genomes means that these instances will be extremely rare because each internal branch is expected to be extremely short. Therefore, it may be that approaches based on maximum parsimony (MP) are sufficiently accurate for reconstructing phylogenies of SARS-CoV-2, and their simplicity means that they can be applied to much larger datasets. Here, we evaluate the performance of *de novo* and online phylogenetic approaches, and ML and MP frameworks, for inferring large and dense SARS-CoV-2 phylogenies. Overall, we find that online phylogenetics produces similar phylogenetic trees to *de novo* analyses for SARS-CoV-2, and that MP optimizations produce more accurate SARS-CoV-2 phylogenies than do ML optimizations. Since MP is thousands of times faster than presently available implementations of ML and online phylogenetics is faster than *de novo*, we therefore propose that, in the context of comprehensive genomic epidemiology of SARS-CoV-2, MP online phylogenetics approaches should be favored.

## Introduction

The widespread availability and extreme abundance of pathogen genome sequencing has made phylogenetics central to combatting the pandemic. Communities worldwide have begun implementing genomic surveillance by systematically sequencing the genomes of a percentage of local cases (Deng et al. 2020; Lu et al. 2020a; Meredith et al. 2020; Park et al. 2021). This has been invaluable in tracing local transmission chains (Bluhm et al. 2020; Lam 2020), understanding the genetic makeup of viral populations within local communities (Gonzalez-Reiche et al. 2020; Franceschi et al. 2021; Thornlow et al. 2021a), uncovering the means by which viral lineages have been introduced to new areas (Castillo et al. 2020), and measuring the relative spread of specific variants (Skidmore et al. 2021; Umair et al. 2021). Phylogenetic approaches for better understanding the proximate evolutionary origins of the virus (Li et al. 2020), as well as to identify recombination events (Jackson et al. 2021; Turakhia et al. 2021b) and instances of convergent evolution (Kalantar et al. 2020; Peng et al. 2021) have greatly informed our understanding of the virus. Phylogenetic visualization software including Auspice (Hadfield et al. 2018) and Taxonium (Sanderson 2021a) have also become widely used for public health purposes.

A comprehensive, up-to-date phylogenetic tree of SARS-CoV-2 is important for public health officials and researchers. A tree containing all available sequences can sometimes facilitate identification of epidemiological links between samples that might otherwise be obscured in subsampled phylogenies. Conversely, these approaches can often rule out otherwise plausible transmission histories. Such information can also help to identify the likely sources of new viral strains in a given area (Moreno et al. 2020; Tang et al. 2021). Additionally, using up-to-date information enables us to find and track quickly growing clades and novel variants of concern (Annavajhala et al. 2021; Tegally et al. 2021), as well as to measure the spread of known variants at both global and community scales. Furthermore, comprehensive phylogenies can facilitate identification of recombinant viral genomes (Turakhia et al. 2021b), natural selection at homoplasious positions (van Dorp et al. 2020), variation in mutation rates (De Maio et al. 2021a), and systematic recurrent errors (Turakhia et al. 2020). This also facilitates naming lineages of interest, which has been especially important in tracking variants of concern during the pandemic (*e.g*. B.1.1.7 or “Alpha” and B.1.617.2 or “Delta”) (Rambaut et al. 2020).

SARS-CoV-2 presents a unique set of phylogenetic challenges. First, the unprecedented pace and scale of whole-genome sequence data has forced the phylogenetics community to place runtime and scalability at the center of every inference strategy. More than 10 million SARS-CoV-2 genome sequences are currently available, with tens of thousands being added each day. Prior to the pandemic, *de novo* phylogenetics, or approaches that infer phylogenies from scratch, have been the standard, as there has rarely been a need to re-infer or improve pre-existing phylogenies on a daily basis. Re-inferring a tree of more than 10 million samples daily, however, is extremely costly, and has brought a renewed focus on methods for adding new samples to existing phylogenetic trees (Matsen et al. 2010; Berger et al. 2011; Izquierdo-Carrasco et al. 2014; Fourment et al. 2018; Barbera et al. 2019). This approach has been called “online phylogenetics” (Gill et al. 2020), and has important advantages in the context of the pandemic and beyond. Online phylogenetics is appealing for the genomic surveillance of any pathogen, because iterative optimization should decrease computational expense, allowing good estimates of phylogenies to be made readily available.

Second, SARS-CoV-2 genomes are much more closely related than sequences in most other phylogenetic analyses. Because the advantages of maximum likelihood methods decrease for closely related samples and long branches are relatively rare in the densely sampled SARS-CoV-2 phylogeny (Felsenstein 1978; Hendy and Penny 1989; Philippe et al. 2005), this suggests that phylogenetic inferences based on maximum parsimony, a much faster and simpler phylogenetic inference method, could be better suited for online phylogenetic analyses of SARS-CoV-2 genomes (Wertheim et al. 2021). The principle of maximum parsimony is that the tree with the fewest mutations should be favored, and it is sometimes described as a non-parametric phylogenetic inference method (Sullivan and Swofford 2001; Kolaczkowski and Thornton 2004). Additionally, because parsimony-based tree optimization does not require estimation of ancestral character state uncertainty at all positions in the phylogeny like ML optimization does, parsimony uses much less memory.

Here, we evaluate approaches that would enable one to maintain a fully up-to-date and comprehensive global phylogeny of SARS-CoV-2 genome sequences (McBroome et al. 2021). Specifically, we investigate tradeoffs between online and *de novo* phylogenetics and between maximum parsimony and maximum likelihood approaches when the aim is for an analysis to scale to millions of sequences, with tens of thousands of new sequences being added daily. We chose to compare maximum parsimony and maximum likelihood (and omit other approaches like neighbor-joining) because they were the most effective methods at inferring large SARS-CoV-2 phylogenies based on previous analyses (Lanfear and Mansfield 2020), and because the most efficient distance-based methods are quadratic in memory usage so cannot scale to estimating trees from datasets of more than a few hundred thousand sequences (Wang et al. 2022). We mimic the time-course of the pandemic by introducing increasingly large numbers of SARS-CoV-2 genome sequences proportionately to their reported sampling dates.

We evaluate potential online phylogenetics approaches by iteratively adding samples to existing trees and optimizing the augmented phylogeny with different tools that have been proposed for this purpose during the pandemic. In particular, we evaluate matOptimize, IQ-TREE 2, and FastTree 2. Between each optimization step, we use UShER (Turakhia et al. 2021a) to add samples to trees by maximum parsimony. matOptimize is a parsimony optimization approach that uses subtree pruning and regrafting (SPR) moves to minimize the total mutations in the final tree topology (Ye et al. 2022). IQ-TREE 2 uses nearest neighbor interchange (NNI) to find the tree with the highest likelihood given an input multiple sequence alignment (Minh et al. 2020). FastTree 2 uses a pseudo-likelihood approach that employs minimum-evolution SPR and/or NNI moves and maximum-likelihood NNI moves while using several heuristics to reduce the search space (Price et al. 2010). The likelihood-based approaches evaluated here report branch lengths in substitutions per site. Parsimony-based matOptimize reports branch lengths in total substitutions, which can be converted to the latter by dividing by the genome length. These branch lengths may be interpreted as is or used as an initial estimate for other distance measures, for example in the construction of time trees (Sanderson 2021b).

Results from our comparisons demonstrate that for the purposes of SARS-CoV-2 phylogenetics, in which samples are numerous and closely related and inference speed is of high significance, parsimony-based online phylogenetics applications are clearly most favorable and are also the only immediately available methods capable of producing daily phylogenetic estimates of all available SARS-CoV-2 genomes (Turakhia et al. 2021a). We note that matOptimize is used to maintain such a phylogeny comprising over 9 million genomes as of April 2022 (McBroome et al. 2021). As similarly vast datasets will soon be available for many species and pathogens, we expect that online approaches using parsimony or pseudo-likelihood optimization will become increasingly central to phylogenetic inference.

## Results and Discussion

### Online phylogenetics is an alternative to de novo phylogenetics for ongoing studies

The vast majority of phylogenetics during the pandemic has consisted of *de novo* phylogenetics approaches (Hadfield et al. 2018; Li et al. 2020; Lu et al. 2020a, 2020b; Meredith et al. 2020), in which each phylogeny is inferred using only genetic variation data, and without a guide tree (Fig. 1). This strategy for phylogenetic inference has long been the default, as in most instances in the past, data are collected just once for a project, and more relevant data are rarely going to be made available in the near future. This process is well characterized and has been foundational for many phylogenetics studies (Hug et al. 2016; Parks et al. 2018; Lu et al. 2020b), and most phylogenetics software is developed with *de novo* phylogenetics as the primary intended usage.

**Figure 1:**
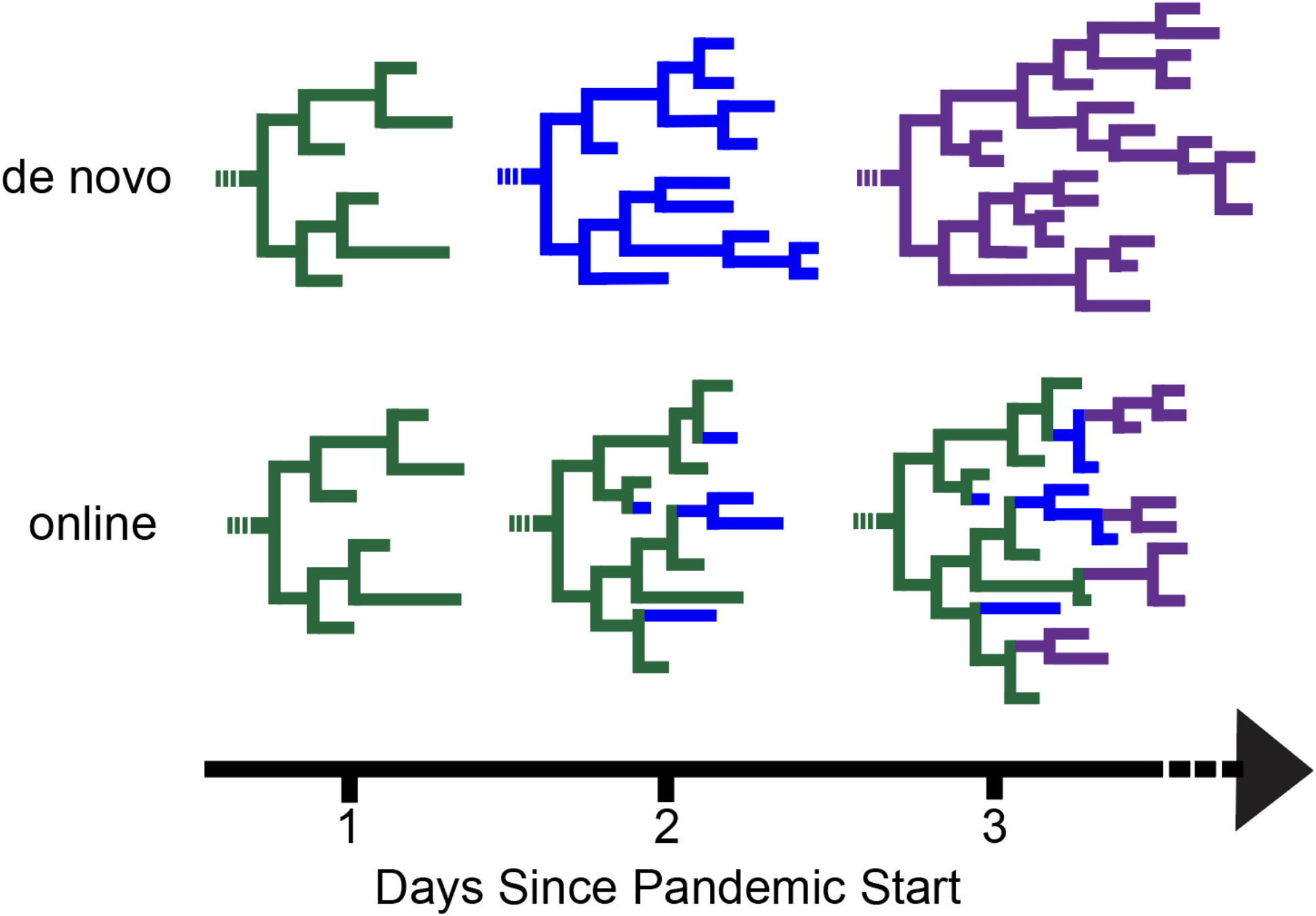
Phylogenies may be optimized from scratch using de novo phylogenetics or iteratively using online phylogenetics. In de novo phylogenetics (top), trees are repeatedly re-inferred from scratch. Conversely, online phylogenetics (bottom) involves placement of new samples as they are collected. Intermittent optimization steps (not depicted) after new samples are placed can help overcome errors from previous iterations. Online phylogenetics is expected to be much faster and require less memory than de novo phylogenetics.

A challenging aspect of pandemic phylogenetics is the need to keep up with the pace of data generation as genome sequences continuously become available. To evaluate phylogenetics applications in the pandemic (Fig. 1), we split 233,326 samples dated from December 23, 2019 through January 11, 2021 into 50 batches according to their date of collection. Each batch contains roughly 5,000 samples. Samples in each batch were collected within a few days of each other, except in the first months of the pandemic when sample collection was more sparse. We also constructed a dataset of otherwise similar data simulated from a known phylogeny (see Methods). The intent of this scheme is to roughly approximate the data generation and deposition that occurred during the pandemic. All datasets are available from the repository associated with this project (Thornlow et al. 2021b), for reproducibility and so that future methods developers can directly compare their outputs to our results. We performed online and *de novo* phylogenetics using a range of inference and optimization approaches. Since thousands of new sequences are added to public sequence repositories each day, we terminated any phylogenetic inference approaches that took more than 24 hours, because such phylogenies would be obsolete for some public health applications by the time they were inferred.

### Analyses using simulated data suggest that online phylogenetics is more accurate for SARS-CoV-2

We first compared matOptimize (commit 66ca5ff, conda version 0.4.8) (Ye et al. 2022), IQ-TREE 2 (Minh et al. 2020), and FastTree 2 (Price et al. 2010) using both online and *de novo* phylogenetics strategies using simulated data that we designed to closely mimic real SARS-CoV-2 datasets. All online phylogenomics workflows used UShER (Turakhia et al. 2021a) to add new sequences to the previous tree (see Methods) as to our knowledge it is the only software package that is fast enough to perform under real time constraints. We chose these three tools based on their widespread usage among SARS-CoV-2 phylogenetics applications (*e.g*. matOptimize is part of the UShER suite (Turakhia et al. 2021a), IQ-TREE 2 is used by (COVID-19 Genomics UK (COG-UK) Consortium 2020; Lanfear and Mansfield 2020) and FastTree 2 is used by (Hadfield et al. 2018)) as well as to cover several different methodologies.

Simulating an alignment based on a known tree ensures that there is a ground truth for comparison to definitively assess each optimization method. We used an inferred global phylogeny as a template to simulate a complete multiple sequence alignment using phastSim (De Maio et al. 2021b). We subsampled this simulated alignment into 50 progressively larger sets of samples, ranging in number of samples from 4,676 to 233,326 (see Methods), to examine each of the three optimization methods in both online and *de novo* phylogenetics. We then computed the Robinson-Foulds distance for unrooted trees of each iteration, after condensing identical samples and collapsing very short branches, to the global mutation-annotated tree on which the simulation was based, pruned to contain only the relevant samples, and normalized by the maximum possible Robinson-Foulds distance between the trees (Fig. 2, Fig. S3) (Steel and Penny 1993).

**Figure 2:**
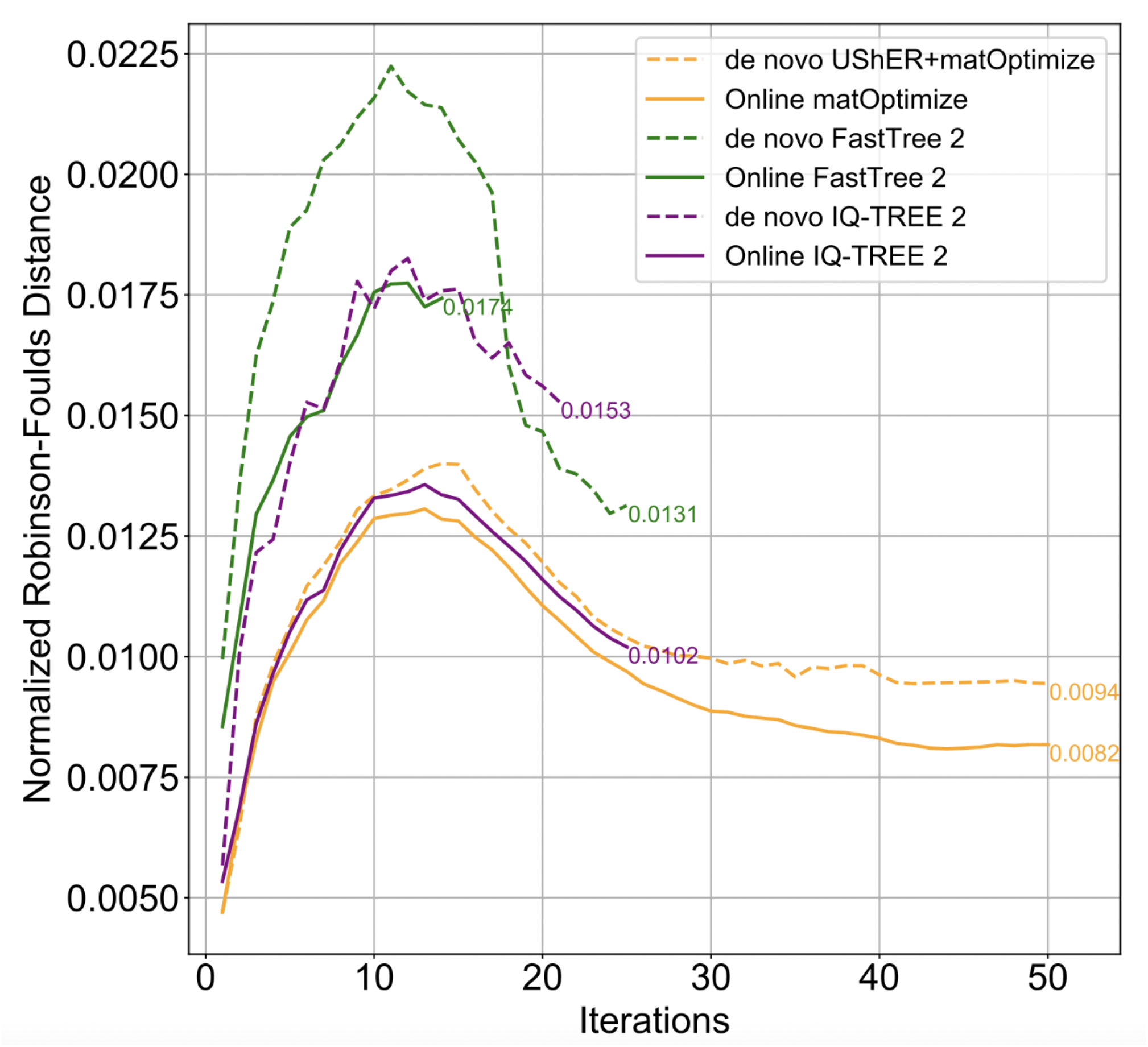
Online matOptimize produces phylogenies most similar to ground truth on simulated data. For each batch of samples, we calculated the Robinson-Foulds distance between the tree produced by a given optimization software and the ground truth tree pruned to contain only the relevant samples. We then normalized these values by the maximum possible Robinson-Foulds distance between the two trees (see Figure S3), which is equal to 2n-6 where n equals the number of samples in each tree (Steel and Penny 1993). We terminated FastTree and IQ-TREE after the first phylogeny that took more than 24 hours to optimize.

All online phylogenetics methods noticeably outperformed their *de novo* counterparts. Overall, online matOptimize produced phylogenies with the lowest Robinson-Foulds distance to the ground truth for the majority of iterations (Fig. 2). Online IQ-TREE 2 performed similarly, but was able to complete only 25 of the 50 iterations due to its extreme computational resource requirements. For example, for the 14th phylogeny of 60,571 sequences, which was the last phylogeny produced using under 200 GB of RAM in under 24 hours by all six methods, we found Robinson-Foulds distances of 1696, 2590, and 2130 for *de novo UShER*+matOptimize, FastTree 2, and IQ-TREE 2 respectively, and distances of 1557, 2111, and 1618 for online matOptimize, FastTree 2, and IQ-TREE 2, respectively.

There are several possible explanations for the improved performance of online phylogenetics relative to *de novo* approaches. First, the radius for SPR moves when optimizing a large tree is insufficiently large to find improvements that are more readily applied when the tree contains fewer samples as in early rounds of online phylogenetics. In online phylogenetics, these improvements carry over to subsequent trees, while in *de novo*, they do not. The radius is defined as the phylogenetic distance of the search space when moving a node to a more optimal position. As the phylogeny increases in size, the distance from a node to its optimal position is likely to also increase, necessitating a larger SPR move radius to make equivalent improvements in larger trees. Second, large clades consisting primarily of samples with branch length zero might further reduce the ability of optimization methods to find improvements by indirectly limiting search space due to the increased number of edges when represented internally as a bifurcating tree. It may sometimes be possible to explore moves across such tree regions during online phylogenetics in early iterations when the polytomy is relatively small. Third, online phylogenetics facilitates tree optimization by providing an exceptionally good initial tree that has already been heavily optimized in previous iterations. We expect that this approach will typically outperform parsimony and neighbor-joining initial trees that are used in most *de novo* phylogenetic inference approaches. Finally, because each online experiment began with a small tree inferred *de novo* by stepwise sample addition with UShER, it is possible that these initial trees are more optimal than initial trees produced during *de novo* inference by the other software we evaluated, perhaps because UShER prefers the reference nucleotide in cases of ambiguous internal character states.

### Analyses using real data suggest that online phylogenetics is more efficient than *de novo* and produces similarly optimal phylogenies

While analyses using simulated data offer the ability to compare to a known ground truth, assessing the performance of each method on real SARS-CoV-2 data may more accurately reflect practical use of each method. Therefore, we also tested each optimization strategy on 50 progressively larger sets of real SARS-CoV-2 samples and calculated the parsimony score and likelihood of each optimized tree, as well as the run-time and peak RAM usage of each software package used (Fig. 3). To accomplish this, we subsampled our global phylogeny, which was produced using stringent quality control steps (see Methods), as before, to mimic the continuous accumulation of samples over the course of the pandemic.

**Figure 3:**
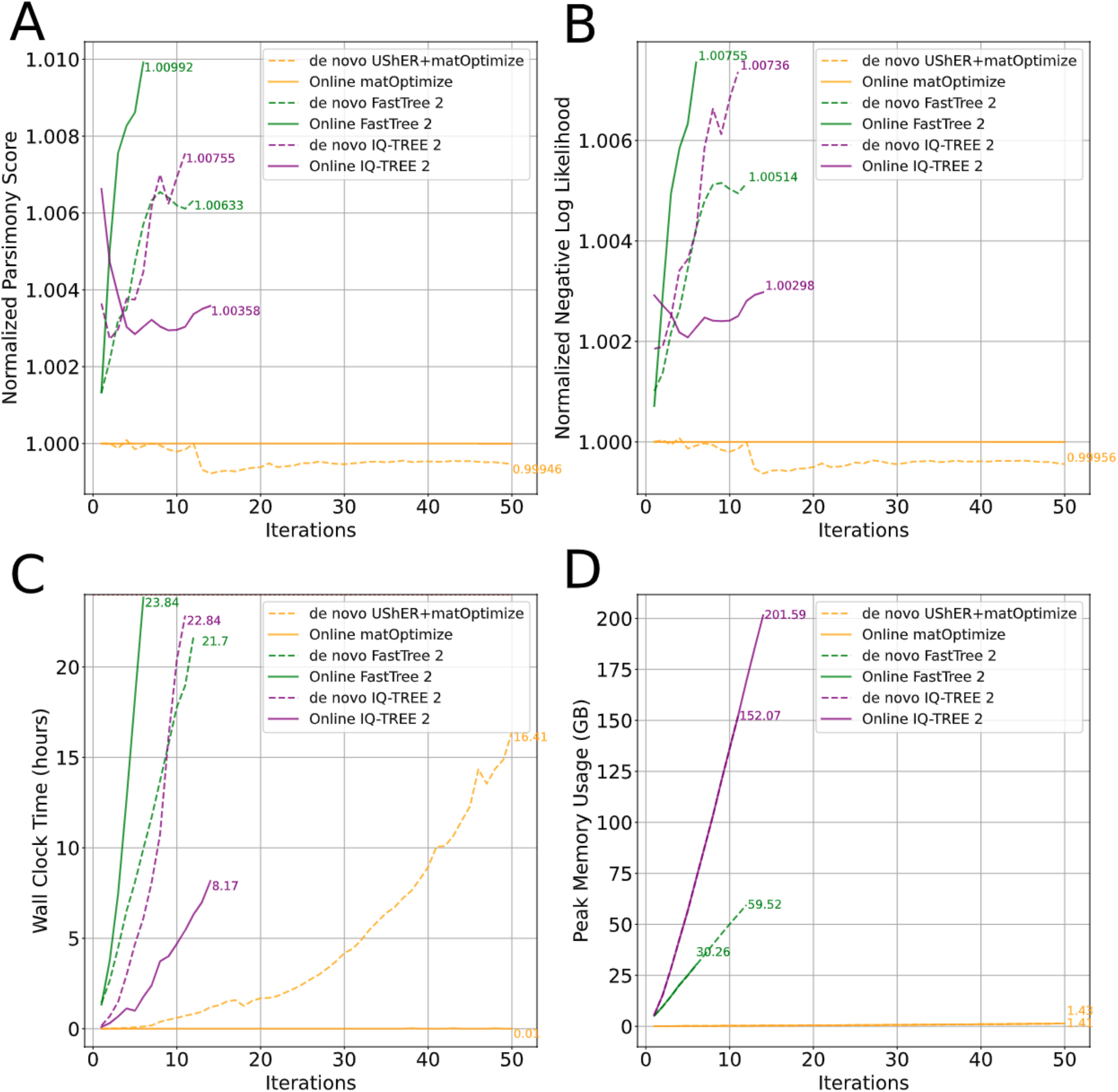
In practice, optimization by parsimony is more effective for SARS-CoV-2 data than optimization by ML. We calculated (A) the parsimony score for each tree using matUtils, (B) the log-likelihood of each tree using IQ-TREE 2, (C) runtime and (D) peak memory usage of each optimization. (A) and (B) are normalized by the value obtained for the matOptimize online approach such that all other methods are expressed as a ratio. Strategies that surpassed 24 hours (C) or the allowable RAM usage (D) were terminated prior. In most cases, with the notable exception of FastTree 2, online phylogenetics (solid lines) perform better than de novo phylogenetics (dashed lines). We ran all matOptimize analyses using an instance with 15 CPUs and 117.2 GB of RAM, and we ran all IQ-TREE 2 and FastTree 2 analyses on an instance with 31 CPUs and 244.1 GB of RAM, but limited each command to 15 threads for equivalence with matOptimize.

Online optimizations are generally much faster than *de novo* phylogenetic inference. For example, IQ-TREE 2 achieves a roughly four-fold faster run-time for online optimizations compared to inferring the tree *de novo* (Fig. 3c). The 11th iteration, which has 47,819 sequences and was the last to be completed by both online and *de novo* IQ-TREE 2, took 22 hours 50 minutes for *de novo* IQ-TREE 2 but only 5 hours 26 minutes for online IQ-TREE 2. *De novo* UShER+matOptimize was the only *de novo* method to finish all trees in fewer than 24 hours, but its speed for each daily update pales in comparison to online matOptimize. Online matOptimize is several orders of magnitude faster than its *de novo* counterpart, and its optimizations for the largest phylogenies take roughly 30 seconds, while *de novo* tree inference with UShER can take several hours for trees consisting of more than 100,000 samples (Fig. 3c). However, whether a software package is used for online or *de novo* phylogenetics does not strongly affect its peak memory usage.

We also found that online phylogenetics strategies produce trees very similar in both parsimony score and likelihood to their *de novo* counterparts, with differences of less than 1% in all cases (Fig. 3a-b). For example, in the 11th iteration containing 47,819 sequences, online IQ-TREE 2 produces a tree with a parsimony score of 32,005, whereas *de novo* IQ-TREE 2 produces a tree with parsimony score 32,149. Our results suggest that in addition to the computational savings that allow online phylogenetics approaches to continuously stay up-to-date, online phylogenetics approaches also produce trees with similar parsimony scores and likelihoods to their *de novo* counterparts.

### Under pandemic time constraints, parsimony-based optimization methods have favorable metrics compared to ML methods for SARS-CoV-2 phylogenies

In the case of both *de novo* and online phylogenetics, the parsimony-based matOptimize outperforms both FastTree 2 and IQ-TREE 2 in runtime and peak memory usage. For the sixth iteration (26,486 samples), which was the largest phylogeny inferred by all online methods in under 24 hours and using under 200 GB of RAM, online FastTree 2 required nearly 24 hours and 30.3 GB of RAM, and online IQ-TREE 2 required 1 hour 45 minutes and 72 GB of RAM. By contrast, matOptimize used only 6 seconds and 0.15 GB of RAM. This iteration contained roughly 10% as many samples as the 50th and final iteration (233,326 total samples), which online matOptimize completed in 32 seconds using 1.41 GB of RAM at peak usage. Even this largest tree represents only a very small fraction of the more than 10 million currently available SARS-CoV-2 genomes, indicating that, among the approaches we evaluated, matOptimize is the only viable option for maintaining a comprehensive SARS-CoV-2 phylogeny via online phylogenetics.

In addition to its scalability, matOptimize outperforms ML optimization methods under 24-hour time constraints in both the parsimony and likelihood scores of the trees that it infers. For the sixth iteration (26,486 samples), we found parsimony scores of 16,130, 16,179, and 16,290 for online matOptimize, IQ-TREE 2, and FastTree 2 respectively. While all methods produce phylogenies with parsimony scores within 1% of each other, matOptimize is consistently the lowest. However, matOptimize was developed to optimize by parsimony, while the other methods were developed for ML optimizations. Unexpectedly, we found log-likelihood scores of −233,414.277, −233,945.528, and −235,177.396 for matOptimize, IQ-TREE 2, and FastTree 2 respectively, indicating that matOptimize produces preferable phylogenies based on likelihood as well. We used a Jukes-Cantor (JC) model to calculate likelihoods due to time constraints in calculation for more complex substitution models, but a Generalised Time Reversible (GTR) model with specified rate parameters produced strongly correlated likelihoods (Fig. S1). Specifically, we fit a generalized linear model using a Gamma family (inverse link function) to predict the likelihood of the tree under the JC model using the iteration of tree construction and the GTR likelihood as predictors. We examined the six trees from the first and second iteration (12 in total). We found that the GTR likelihood was significantly correlated with the JC likelihood (p < 2.27×10^-5^).

### Parsimony optimization produces comparable or more favorable SARS-CoV-2 trees than the most thorough maximum likelihood methods

We also compared the performance of *de novo* inference with UShER+matOptimize to state-of-the-art methods without a 24-hour limit on runtime. In three iterations of increasing size (~4.5k, ~8.9k, and ~13.2k samples), we inferred trees from real and simulated data using UShER+matOptimize, IQ-TREE 2 with stochastic search enabled, and RAxML-NG. With the parameters used here, IQ-TREE 2 performs stochastic NNI moves in addition to hill-climbing NNI. RAxML-NG is a maximum likelihood approach that uses SPR moves to search tree-space for higher likelihood phylogenies (Kozlov et al. 2019). We allowed each experiment to run for up to two weeks. All programs completed successfully on the first iteration. RAxML-NG did not terminate within two weeks for the second and third iterations. On real data, we found that UShER+matOptimize produced trees with higher log-likelihoods than IQ-TREE 2 and RAxML-NG across all three iterations (Fig. 4A). Under the substitution model parameters estimated by IQ-TREE 2, the log-likelihoods for the first iteration were −73780.756, −73828.271, and −73782.289 for UShER+matOptimize, IQ-TREE 2, and RAxML-NG respectively. Under the parameters estimated by RAxML-NG, the log-likelihoods for the first iteration were −73754.894, −73801.935, and −73756.246 for UShER+matOptimize, IQ-TREE 2, and RAxML-NG respectively. On simulated data, UShER+matOptimize produced trees closer to the ground truth than the other methods when measured by quartet distance across all three iterations (Fig. 4B). By RF distance, the UShER+matOptimize trees were closest to the ground truth for the second and third iterations, but the RAxML-NG tree was closest to ground truth in the first iteration (Fig. 4C). We therefore conclude that parsimony-based tree inference can perform equivalently or better than state of the art maximum likelihood approaches but do this in a tiny fraction of the time, making it by far the most suitable approach for pandemic-scale phylogenetics of SARS-CoV-2.

**Figure 4:**
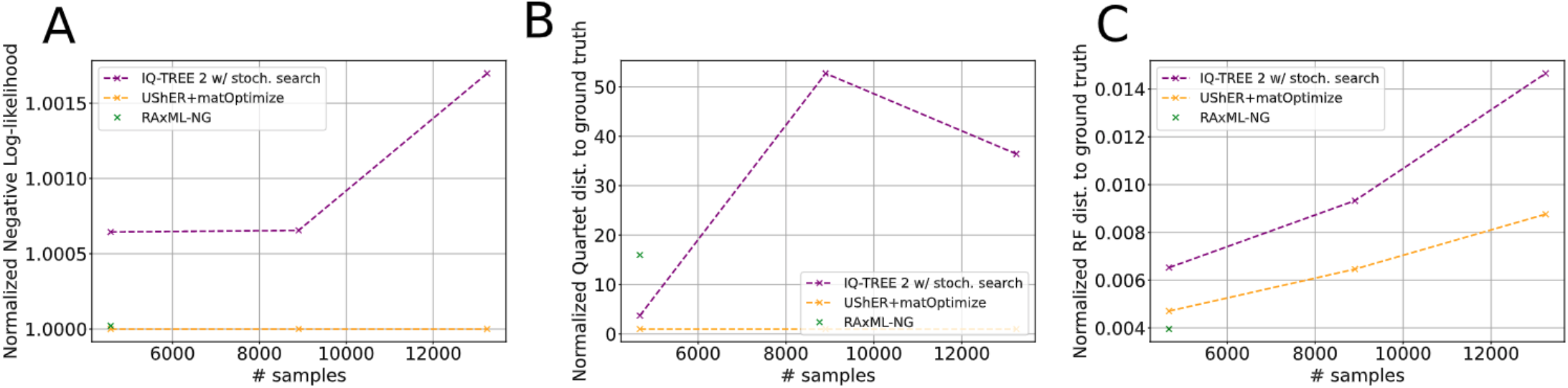
de novo matOptimize produces similar or more favorable trees compared to the most thorough maximum-likelihood inference programs. We ran these methods for up to two weeks each to infer trees de novo from the three smallest iterations of real and simulated data. For real data **(A),** log-likelihoods were computed under the model parameters estimated by IQ-TREE 2 at each iteration. Values are normalized by the value of the matOptimize approach such that all other methods are expressed as a ratio. For simulated data **(B, C)**, the reported quartet distances **(B)** are similarly normalized by the value of the matOptimize approach such that other methods are expressed as a ratio. RF distances **(C)** are normalized by the maximum possible RF distance of 2n-6, where n is the number of leaves in each tree. For all panels, the second and third iterations of RAxML-NG (which did not terminate within two weeks) are omitted.

### Parsimony and likelihood are strongly correlated when optimizing large SARS-CoV-2 phylogenies

While our comparisons of online and *de novo* as well as parsimony-based and ML optimizations of cumulative pandemic-style data demonstrated practical performance, the largest trees completed by all methods in these experiments represent only a small fraction of available SARS-CoV-2 data. It is also crucial that we identify the optimal ways to produce a large phylogeny from already aggregated data. We therefore evaluated phylogenetic inference methods for optimizing a tree of 364,427 SARS-CoV-2 genome sequences, without constraining methods according to time or memory requirements. We optimized this global phylogeny using matOptimize (Ye et al. 2022), IQ-TREE 2 (Minh et al. 2020), and FastTree 2 (Price et al. 2010). Overall, we found that matOptimize produced the tree with the lowest parsimony score across all methods in roughly one hour (Table 1).

**Table 1:** We applied each of the three optimization methods to a starting tree of 364,427 SARS-CoV-2 samples, which had an initial parsimony score of 296,247. We first ran 2 iterations of IQ-TREE 2 optimization, using an SPR radius of 20 on the first and 100 on the second. We also used an SPR radius of 10 on one iteration of matOptimize, and six iterations of pseudo-likelihood optimization using FastTree 2, which we terminated after roughly 10.5 days.

| Method | Iterations | Runtime (H:M:S) | Final Parsimony Score (Percent Change from Starting Tree) |
| --- | --- | --- | --- |
| IQ-TREE 2 | 2 | 24:30:52 | 294,258 (0.67) |
| FastTree 2 | 6 | 252:02:49 | 294,154 (0.71) |
| matOptimize | 1 | 1:12:03 | 294,022 (0.75) |

We found that after each of the six iterations of FastTree 2 optimization, the likelihood and parsimony improvements are strongly linearly correlated (Fig. 5). This suggests that changes achieved by maximizing parsimony will also optimize likelihood for SARS-CoV-2 data. That is, for extremely densely sampled phylogenies wherein long branches are especially rare, parsimony and likelihood of phylogenies, and tree moves to optimize either are highly correlated. However, despite the strength of this correlation, we find an extreme disparity in practical usage when optimizing by either metric. Parsimony-based methods are far more time- and data-efficient, and presently available ML approaches quickly become prohibitively expensive. For example, while the 6 iterations of FastTree did result in large improvements in both likelihood and parsimony score, the resulting tree would be out of date long before the 10.5-day optimization had completed. Moreover, we applied matOptimize to the tree output by the sixth iteration of FastTree, achieving a parsimony score of 293,866 (improvement of 288) in just 16 minutes, indicating that even after 10.5 days, additional optimization was still possible. This suggests that, for the purposes of optimizing even moderately large SARS-CoV-2 trees, parsimony-based methods should be heavily favored due to their increased efficiency.

**Figure 5.**
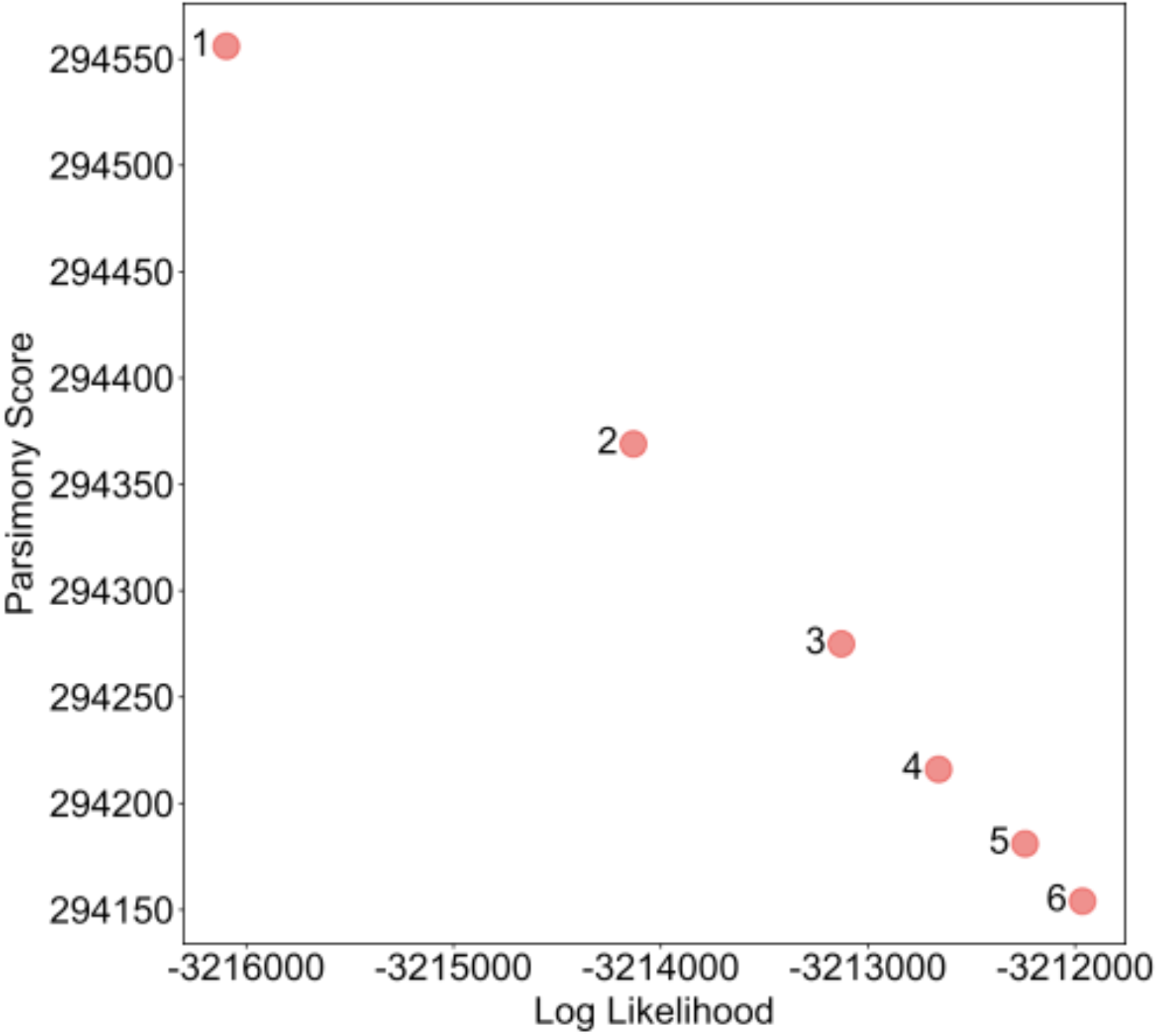
Improvement in likelihood and parsimony have a linear relationship for our optimized global tree. We optimized our initial global tree using 6 iterations of FastTree and measured the total parsimony and the likelihood after each, finding a linear relationship (Pearson correlation, rho = −1.0, p < 2.9×10^-7^).

## Conclusions

The SARS-CoV-2 pandemic has made phylogenetics central to efforts to combat the spread of the virus, but has posed challenges for many commonly used phylogenetics frameworks. A major component of this effort relies on a comprehensive, up-to-date, global phylogeny of SARS-CoV-2 genomes. However, the scale and continuous growth of the data have caused difficulties for standard *de novo* phylogenetic methods. Here, we find that online phylogenetics methods are practical, pragmatic, and accurate for inferring daily phylogenetic trees from a large and densely-sample virus outbreak.

One counterintuitive result is that parsimony-based optimizations outperform sufficiently efficient ML approaches regardless of whether phylogenies are evaluated using parsimony or likelihood. This might be a consequence of the fact that parsimony scores and likelihoods are strongly correlated across phylogenies inferred via a range of phylogenetic approaches. The extremely short branches (Fig. S2) on SARS-CoV-2 phylogenies mean that the probability of multiple mutations occurring at the same site on a single branch is negligible. Stated another way, SARS-CoV-2 is approaching a “limit” where parsimony and likelihood are nearly equivalent. In turn, because of their relative efficiency, parsimony-based methods are able to search more of the possible tree space in the same amount of time, thereby resulting in trees with better likelihoods and lower parsimony scores than trees optimized using currently-available ML software packages. We emphasize that this does not bear on the relative merits of the underlying principles of ML and MP, but instead reflects the utility of methods that have been applied during the pandemic. Nevertheless, this observation does suggest that in some cases, MP optimization may provide a fast and accurate starting point for ML optimization methods. Indeed, many popular phylogenetics software, such as RAxML (Stamatakis 2014) and IQ-TREE (Minh et al. 2020) already use stepwise-addition parsimony trees as initial trees for their optimization. Our results suggest that further optimization of these initial trees using MP may provide benefits in speed *and* accuracy for some datasets, even when the target is an estimate of the ML tree.

As sequencing technologies progress and become more readily available, sample sizes for phylogenetic analyses of major pathogens and highly-studied organisms will necessarily continue to increase. Today, SARS-CoV-2 represents an extreme with respect to the total number of samples relative to the very short branch lengths on the phylogeny. However, the global sequencing effort during the pandemic suggests that the public health sphere has a strong interest in the increased application of whole-genome sequencing to study the genomic contents, evolution, and transmission history of major and emerging human pathogens. We expect that million-sample datasets will become commonplace in the near future. Parsimony-based methods like matOptimize show promise for huge datasets with short branch lengths. Similarly, recently developed parsimony-based likelihood approximations may ultimately be similarly scalable and accurate (De Maio et al. 2022). Online phylogenetics using both of these methods will be a fruitful avenue for future development and application to accommodate these datasets.

## Methods

We first developed a “global phylogeny”, from which all analyses in this study were performed. We began by downloading VCF and FASTA files corresponding to March 18, 2021 from our own daily-updated database (McBroome et al. 2021). The VCF file contains pairwise alignments of each of the 434,063 samples to the SARS-CoV-2 reference genome. We then implemented filters, retaining only sequences containing at least 28,000 non-N nucleotides, and fewer than two non-[ACGTN-] characters. We used UShER to create a phylogeny from scratch using only the remaining 366,492 samples. To remove potentially erroneous sequences, we iteratively pruned this tree of highly divergent internal branches with branch parsimony scores greater than 30, then terminal branches with branch parsimony scores greater than 6, until convergence, resulting in a final global phylogeny containing 364,427 samples. The branch parsimony score indicates the total number of substitutions along a branch. Similar filters based on sequence divergence are used by existing SARS-CoV-2 phylogenetic inference methods. For full reproducibility, files used for creating the global phylogeny can be found in subrepository 1 on the project GitHub page (Thornlow et al. 2021b).

Following this, we tested several optimization strategies on this global phylogeny, hereafter the “starting tree”. We used matOptimize, FastTree 2, and maximum parsimony (MP) IQ-TREE 2. MP IQ-TREE 2 uses parsimony as the optimality criterion in contrast to the maximum likelihood mode used in all other experiments, which was infeasible on a dataset of this size. In these optimization experiments, we used experimental versions of MP IQ-TREE 2 that allow finer control of parsimony parameters (specific versions are listed in the supplemental Github repository). In one experiment, we used the starting tree and its corresponding alignment and ran five iterations of MP IQ-TREE 2, varying the SPR radius from 20 to 100 in increments of 20. Experiments on a small dataset indicated that there is little or no improvement in parsimony score beyond a radius of 100. Separately, we tested another strategy that applied two iterations of MP IQ-TREE 2 to the starting tree, the first iteration using an SPR radius of 20 and the second using a radius of 100. Finally, we tested a strategy of six iterations of pseudo-likelihood optimization with FastTree 2 followed by two iterations of parsimony optimization with matOptimize. The tree produced by this strategy, hereafter the “ground truth” tree, had the highest likelihood of all the strategies we tested. This tree (after_usher_optimized_fasttree_iter6.tree) and files for these optimization experiments can be found in subrepository 2.

In the multifurcating ground truth tree of 364,427 samples, there are 265,289 unique (in FASTA sequence) samples. There are 447,643 nodes in the tree. For reference, a full binary tree with the same number of leaves has 728,853 nodes. 23,437 of the 29,903 sites in the alignment are polymorphic (they display at least two non-ambiguous nucleotides). Homoplasies are common in these data. In the starting tree, 19,090 sites display a mutation occurring on at least two different branches, and 4,976 sites display a mutation occurring more than ten times in the tree.

To mimic pandemic-style phylogenetics, we separated a total of 233,326 samples from the starting tree of 364,427 samples into 50 batches of ~5,000 by sorting according to the date of sample collection. We then set up two frameworks for each of the three software packages (matOptimize (commit 66ca5ff, conda version 0.4.8), maximum-likelihood IQ-TREE 2 (multicore version 2.1.3 COVID-edition), and FastTree 2 (Double Precision version 2.1.10)). The online phylogenetics frameworks began by using UShER to infer a small tree *de novo* from the first batch of samples, followed by alternating steps of optimization using one of the three evaluated methods and placement of additional samples with UShER. In *de novo* phylogenetics, we supplied each software package with an alignment corresponding to all samples in that batch and its predecessors (or VCF for matOptimize) without a guide tree. For both cases, each tree is larger than its predecessor by ~5,000 samples, and each tree necessarily contains all samples in the immediately preceding tree. For FastTree 2, we used 2 rounds of subtree-prune-regraft (SPR) moves (-spr 2), maximum SPR length of 1000 (-sprlength 1000), zero rounds of minimum evolution nearest neighbor interchanges (-nni 0), and the Generalised Time Reversible + Gamma (GTR+G) substitution model (-gtr -gamma). For IQ-TREE 2, we used a branch length minimum of 0.000000001 (-blmin 1e-9), zero rounds of stochastic tree search (-n 0), and the GTR+G substitution model (-m GTR+G). With these parameters, IQ-TREE 2 constructs a starting parsimony tree and then performs hill-climbing NNI steps to optimize likelihood, avoiding the significant time overhead of stochastic search. We ran all matOptimize analyses using an instance with 15 CPUs and 117.2 GB of RAM, and we ran all IQ-TREE 2 and FastTree 2 analyses on an instance with 31 CPUs and 244.1 GB of RAM, but we limited each command to 15 threads for equivalence with matOptimize. Files for all simulated data experiments can be found in subrepository 3.

To generate our simulated data, we used the SARS-CoV-2 reference genome (GISAID ID: EPI_ISL_402125; GenBank ID: MN908947.3) (Shu and McCauley 2017; Sayers et al. 2021) as the root sequence and used phastSim (De Maio et al. 2021b) to simulate according to the ground truth phylogeny described above. Intergenic regions were evolved using phastSim using the default neutral mutation rates estimated in ref. (De Maio et al. 2021a), with position-specific mean mutation rates sampled from a gamma distribution with alpha=beta=4, and with 1% of the genome having a 10-fold increase mutation rate for one specific mutation type (SARS-CoV-2 hypermutability model described in ref. (De Maio et al. 2021b)). Evolution of coding regions was simulated with the same neutral mutational distribution, with a mean nonsynonymous/synonymous rate ratio of omega=0.48 as estimated in (Turakhia et al. 2021a), with codon-specific omega values sampled from a gamma distribution with alpha=0.96 and beta=2. Rates for each intergenic and coding region were not normalized in order to have the same baseline neutral mutation rate distribution across the genome.

We repeated our iterative experiments using *de novo* and online matOptimize, IQ-TREE 2 and FastTree 2 on this simulated alignment, using the same strategies as before. However, instead of computing parsimony and likelihood scores, we computed the Robinson-Foulds (RF) distance (Robinson and Foulds 1981) of each optimization to the ground truth tree, pruned to contain only the samples belonging to that batch. To calculate each RF distance, we used the -O (collapse tree) argument in matUtils extract (McBroome et al. 2021) and then used the dist.topo command in the *ape* package in R (Paradis and Schliep 2019), comparing the collapsed optimized tree and the pruned, collapsed ground truth tree at each iteration. We computed normalized RF distances as a proportion of the total possible RF distance, which is equivalent to two times the number of samples in the trees minus six (Steel and Penny 1993).

Eliminating the 24-hour runtime restriction, we also repeated the first three *de novo* iterative experiments on both real and simulated data to compare UShER+matOptimize, IQ-TREE 2 with stochastic search, and RAxML-NG. These iterations of ~4.5k, ~8.9k, and ~13.2k samples were allowed to run for up to 14 days. For runs that did not terminate within this time (the second and third iterations of RAxML-NG), we used the best tree inferred during the run for comparisons. We ran IQ-TREE 2 and RAxML-NG under the GTR+G model with the smallest minimum branch length parameter that did not cause numerical errors. To compare the trees inferred from real data, we computed log-likelihoods under the GTR+G model for all trees, fixing the model parameters to those estimated by IQ-TREE 2 during tree inference. We also compared the log-likelihoods of the trees under the parameters estimated by RAxML-NG for the first iteration, but could not do so for the second and third iterations which did not terminate in under two weeks. We allowed optimization of branch lengths during likelihood calculation. For the UShER+matOptimize trees, before computing likelihoods, we converted the branch lengths into units of substitutions per site by dividing each branch length by the alignment length (29,903). To compare the trees inferred from simulated data, we computed the RF and quartet distances of each tree to the corresponding ground truth tree described above.

---

**Figure S1:**
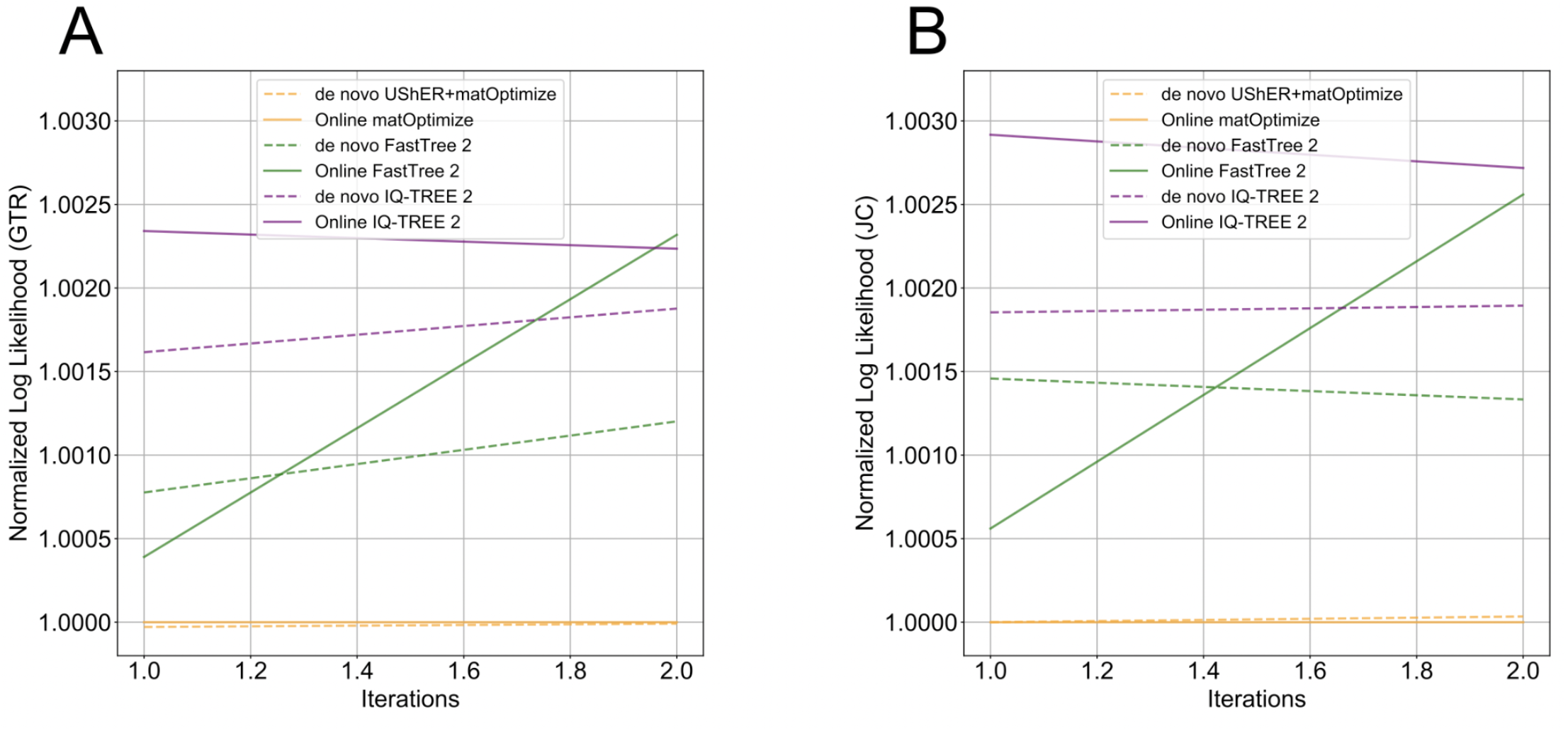
Log-likelihoods calculated using Generalised Time Reversible (GTR) and Jukes-Cantor (JC) models are correlated. We calculated log-likelihoods for each de novo and online method as in Figure 2B using (A) GTR+G and (B) JC models, which suggest that relative performance of each method is consistent across models, and significantly correlated with each other. All values are normalized by the value obtained for the matOptimize online approach, such that other methods are expressed as a ratio.

**Figure S2:**
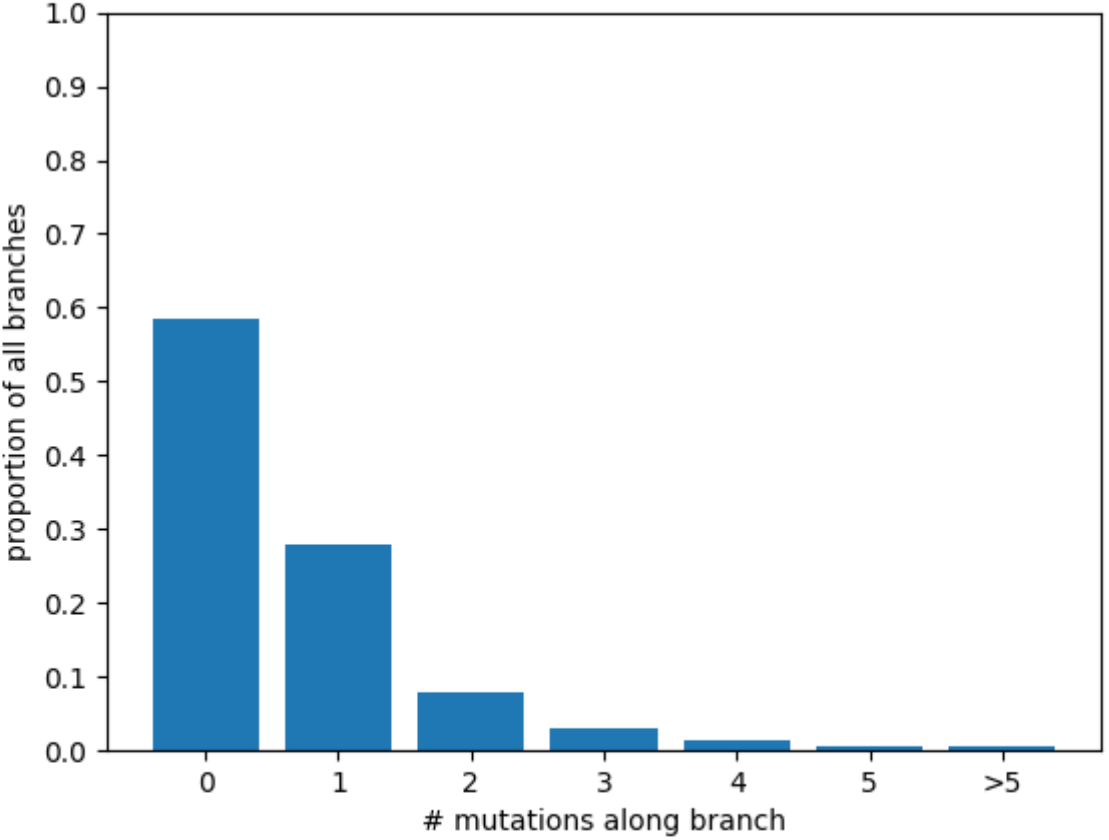
Most branches in the ground truth phylogeny are extremely short. In our optimized global SARS-CoV-2 phylogeny, the majority of branch lengths are zero. This low amount of divergence yields many identical nodes in the tree and demonstrates that the probability of observing multiple mutations at a single site along the same branch is negligible. These characteristics may help explain the ability of parsimony-based inference methods to outperform likelihood optimization on SARS-CoV-2 data.

**Figure S3:**
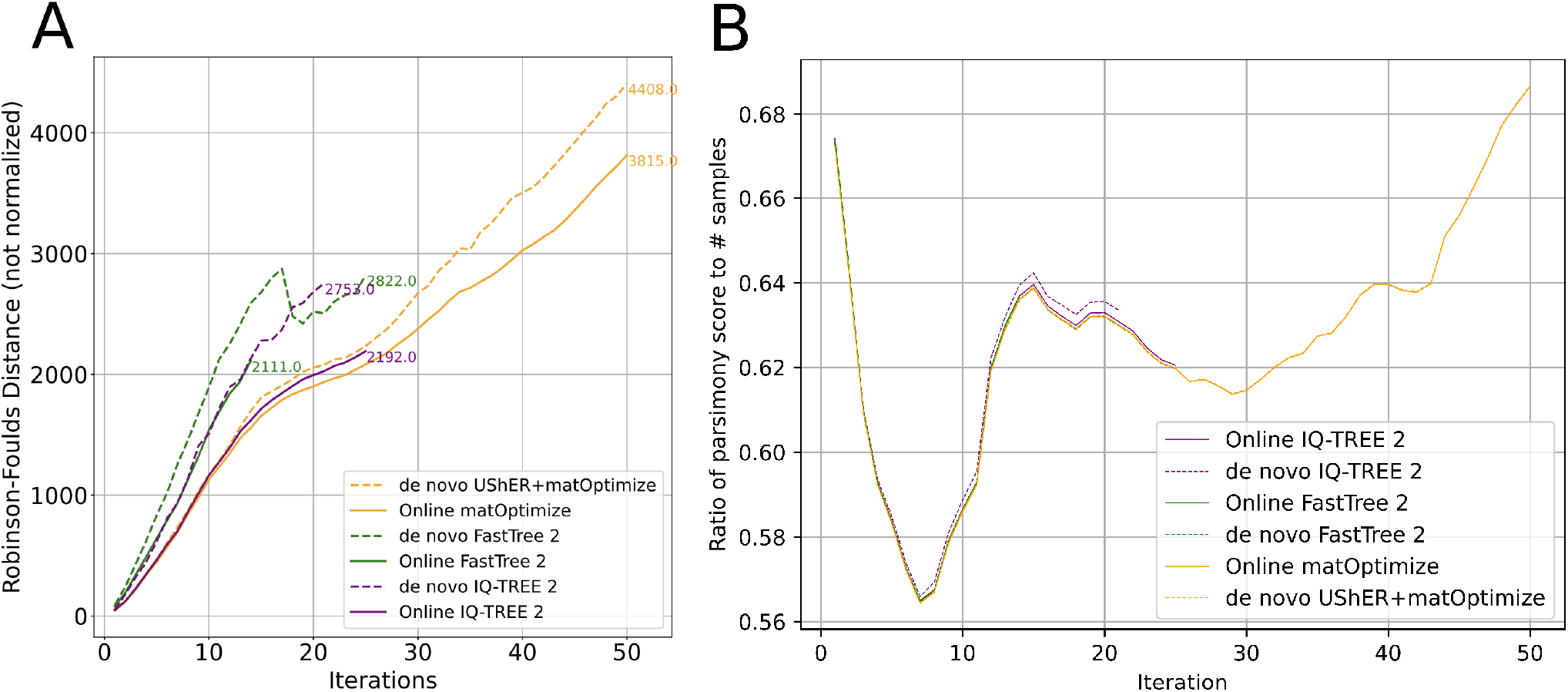
Temporal patterns in simulated SARS-CoV-2 data may affect Robinson-Foulds (RF) distance normalization. The RF distances for each tree in Figure 2 are normalized against the maximum possible RF distance for that tree. While the raw RF distances are approximately continuously increasing **(A)**, they do not increase linearly with the maximum RF distance, leading to the pattern observed in Figure 2. A potential explanation for this is the variation in sequence diversity over simulated time. The ratio of the number of total mutations in the tree (parsimony score) to the number of samples in the inferred trees at each iteration **(B)** approximates the average divergence between samples in each tree. The initial drop in divergence per sample may contribute to the more rapid increase in RF distance because there is less phylogenetic signal to facilitate the resolution of correct topologies. As the divergence subsequently increases, tree inference improves before the RF distances stabilize and begin to increase approximately linearly.

## Acknowledgments

We gratefully acknowledge the authors from the originating laboratories responsible for obtaining each sample, as well as the submitting laboratories where the genome data were generated and shared, on which this research is based.

## Funding

This work was supported by National Institutes of Health (R35GM128932 to R.C.D., T32HG008345 (B.T. and J.M.), F31HG010584 to B.T.), Alfred P. Sloan Foundation fellowship, University of California Office of the President Emergency COVID-19 Research Seed Funding (R00RG2456 to R.C.-D.), European Molecular Biology Laboratory (to N.D.M.), Australian Research Council (DP200103151 to R.L.), Chan-Zuckerberg Initiative grant (to R.L.), and by Eric and Wendy Schmidt by recommendation of the Schmidt Futures program.

## Competing interests

R.L. worked as an advisor to GISAID from mid 2020 to late 2021. The remaining authors declare no competing interests.

## Notes

https://github.com/bpt26/parsimony/

### Competing Interest Statement

Robert Lanfear works as an advisor to GISAID. The remaining authors declare no competing interests.

### Summary of Updates

Added Alexander Kramer as co-first author, significantly revised figures and text

